# Directional stepping model for yeast dynein: Longitudinal- and side- step distributions

**DOI:** 10.1101/645796

**Authors:** Itay Fayer, Rony Granek

**Affiliations:** The Stella and Avram Goren-Goldstein Department of Biotechnology Engineering, Ben-Gurion University of the Negev, Beer Sheva 84105, Israel; Ilse Katz Institute for Meso and Nanoscale Science and Technology, Ben-Gurion University of the Negev, Beer Sheva 84105, Israel

## Abstract

We deduce the directional step distribution of yeast dynein motor protein on the microtubule surface by combing intrinsic features of the dynein and microtubule. These include the probability distribution of the separation vector between the two microtubule binding domains (MTBDs), the angular probability distribution of a single MTBD translation, the existence of a microtubule seam defect, microtubule binding sites, and theoretical extension that accounts for a load force on the motor. Our predictions are in excellent accord with the measured longitudinal step size distributions at various load forces. Moreover, we predict the side-step distribution and its dependence on longitudinal load forces, which shows a few surprising features. First, the distribution is broad. Second, in the absence of load, we find a small right-hand bias. Third, the side-step bias is susceptible to the longitudinal load force; it vanishes at a load equal to the motor stalling force and changes to a left-hand bias above that value. Fourth, our results are sensitive to the ability of the motor to explore the seam several times during its walk. While available measurements of side-way distribution are limited, our findings are amenable to experimental check and, moreover, suggest a diversity of results depending on whether the microtubule seam is viable to motor sampling.

**Significance Statement:** The function of microtubule (MT) associated protein motors, kinesin and dynein, is essential for a myriad of intracellular processes. Different measurements on yeast cytoplasmic-dynein stepping characteristics appear to be unrelated to each other. We provide a unified physical-statistical model that combines these seemingly independent features with a theoretical expression that accounts for the exertion of a longitudinal load force, to yield the longitudinal step distribution at various load forces. The latter is in excellent accord with the measured distributions. Moreover, we deduce the side-step distribution, which surprisingly is susceptible to longitudinal load forces and comprises a right or left bias. This side-way bias is consistent with observations of helical motion of a nanoparticle carried by a number of motors.

## Introduction

The function of microtubule (MT) associated protein motors is essential for a myriad of intracellular processes, such as cargo transfer, cell motility, and cell division. Two of the most studied MT-associated motor proteins are cytoplasmic-dynein (or, in short, dynein) and kinesin; these motors move either towards the MT minus-end (dynein) or MT plus-end (kinesin) (1). For intracellular dynein transport, this implies motion towards the centrosome (i.e., the microtubule organization center) that resides in the vicinity of the nucleus, whereas kinesin advances from the cell nucleus towards the cell periphery. In this work, we focus on dynein. Dynein is a homodimer motor protein; each subunit of dynein is composed of a tail, linker, AAA+ ring (“head”), stalk, buttress, and microtubule-binding domain (MTBD) (2) (see Fig. 1). In broad brushstrokes, cytoplasmic dynein motors can be divided into two sub-families: mammalian and yeast dynein, or weak and strong dynein (according to their characteristic stalling force (3, 4)), respectively. There is a burgeoning interest in these motors (2–18), in particular, their mechanism of motion and regulation. However, dynein’s stepping mechanism remains largely unknown. Furthermore, in recent years, evidence emerged that, unlike kinesin, yeast cytoplasmic dynein motors (henceforth referred to as “dynein”) are not limited to steps that are parallel to the MT long axis (*x*-axis), and in fact, they perform steps in all directions, such as in the direction of the MT short axis (*y*-axis) (5–7). While our primary focus in this work is yeast dynein, some conclusions will be drawn regarding mammalian cytoplasmic dynein.

**Fig. 1.**
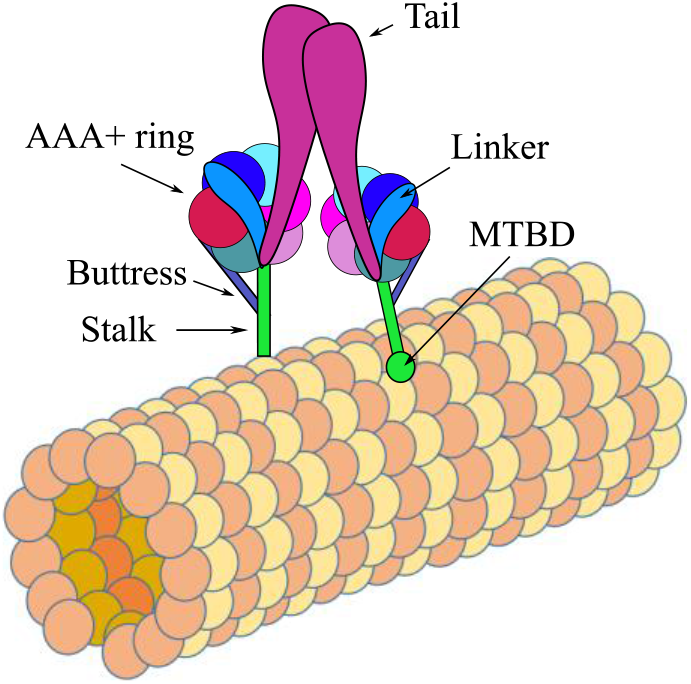
Illustration of dynein’s structure (NOT to scale).

Previous studies showed the existence of dynein *longitudinal* motion towards and away from the MT’s minus end, with and without exertion of external force (3, 8). So far, most studies focused on the *longitudinal* motion, yet, only a few of them (5–7) detected dynein side-steps, presumably due to experimental difficulties. In this work, we deduce the *vectorial* (directional) step distribution of dynein microtubule binding domains (MTBDs) and pivot. Since dynein’s two heavy chains are dimerized *via* their tail domains (2), each MTBD movement depends on the location of the other MTBD of the same dynein, and the dynein pivot movement is dictated by the MTBD translations. Thus, knowledge of the MTBDs *vectorial* (directional) step distribution is crucial for the prediction of dynein pivot motion. Our computation is based on the following key elements: (i) prior knowledge of the binding sites of the MT to the dynein MTBD (19) (20), (ii) the measured probability distribution of the separation vector between the two MTBDs belonging to the same dynein motor (7), (iii) the angular probability distribution of a single MTBD translation (7), and (iv) theoretical extension to the existence of a load (either positive, i.e. pulling, or negative, i.e. assisting) force acting on the motor.

The computed distribution for the vectorial step 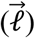 of MTBDs is then split into distributions for *ℓ_x_* (the component of 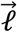 along the *x*-axis), describing *longitudinal* steps, and *ℓ_y_* (the component of 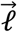 along the *y*-axis), describing side-steps. In a similar manner, the vectorial step of dynein pivot is denoted as 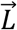, with its components *L_x_* and *L_y_* along the *x* and *y*-axes, respectively. The mechanical balance between the two MTBDs implies that the dynein pivot is always located between the two MTBDs, hence 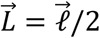, allowing the calculation of the pivot step distribution.

These computed distributions are compared to those measured. In what follows we describe the above model inputs in more detail.

## Methods

### Dynein stepping model

#### 1. Dynein MTBDs only bind to specific binding sites (MT-MTBD sites)

The MT is a hollow cylinder with a radius of ~12.5 nm and composed of ~13 parallel protofilaments that are slightly displaced with respect to one another in the direction of the MT *x*-axis, creating a 3-start pseudo helix structure (21). This pseudo-helical structure is created due to a mismatch that breaks the MT helical continuity, also termed as “lattice seam” or in short “seam” (22). Each protofilament contains repeating and alternating α- and β-tubulin proteins(21). It was demonstrated that MTBD attaches to the MT surface mainly at specific sites that are located between the α- and β-tubulin (19, 20). Thus, we assume here that MTBDs can only attach to these sites. Under this assumption, an examination of the top projection, i.e., “bird’s-eye view,” of an MT with its seam located at the bottom (see Fig. 2-b), reveals a 2D array of attachment sites which resides on *transverse* diagonals (see Fig. 2-c). The *longitudinal* distance between each neighboring pair of diagonals is 8 nm (21), and the angle between each diagonal and the *y*-axis is approximately 0.15 radians (or 7.5 degrees) (see Fig. 3 and Supplementary Information [SI] Fig.1-b). We further assume that, due to obvious steric effects, MTBDs of a dynein – whose pivot is located in the middle of the cylindrical “upper” half – cannot attach to sites that reside on the cylindrical “lower” half (see Fig.2-b & 2-c). Consequently, the width of the 2D array is 25 nm.

**Fig. 2.**
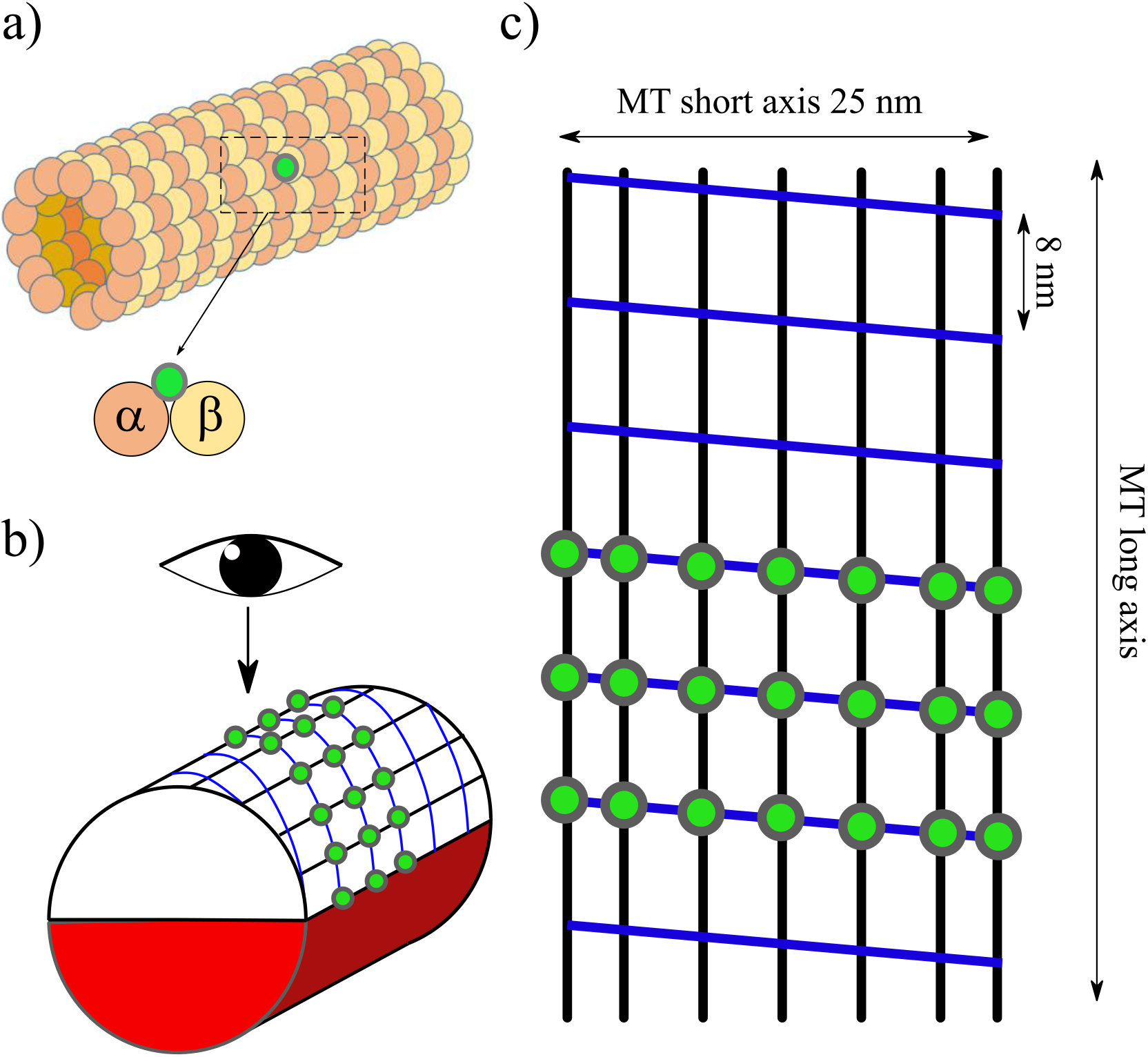
Illustration of the 2D attachment sites array: **(a)** MT 3D structure: alpha- and beta-tubulin are denoted as orange and yellow circles, respectively, and an example of an MT-binding site to the MTBD is denoted as a green dot. **(b)** MT structure as a cylinder with a grid of binding sites. Binding sites are denoted as green dots. The arrow indicates the angular position of a dynein pivot (“center of mass”). The “lower” MT cylindrical half, colored in red, indicates a forbidden area where MTBDs cannot bind due to steric effects. **(c)** The grid of binding sites, or 2D array, as projected from above with respect to dynein pivot/center of mass. The blue lines are the “transverse diagonals” referred to in the main text, and the black lines denote the tubulin protofilaments. The longitudinal distance (along the MT long axis) between each pair of adjacent diagonals is 8 nm, and the width of the 2D array (along the MT short axis) is 25nm, the MT diameter.

**Fig. 3.**
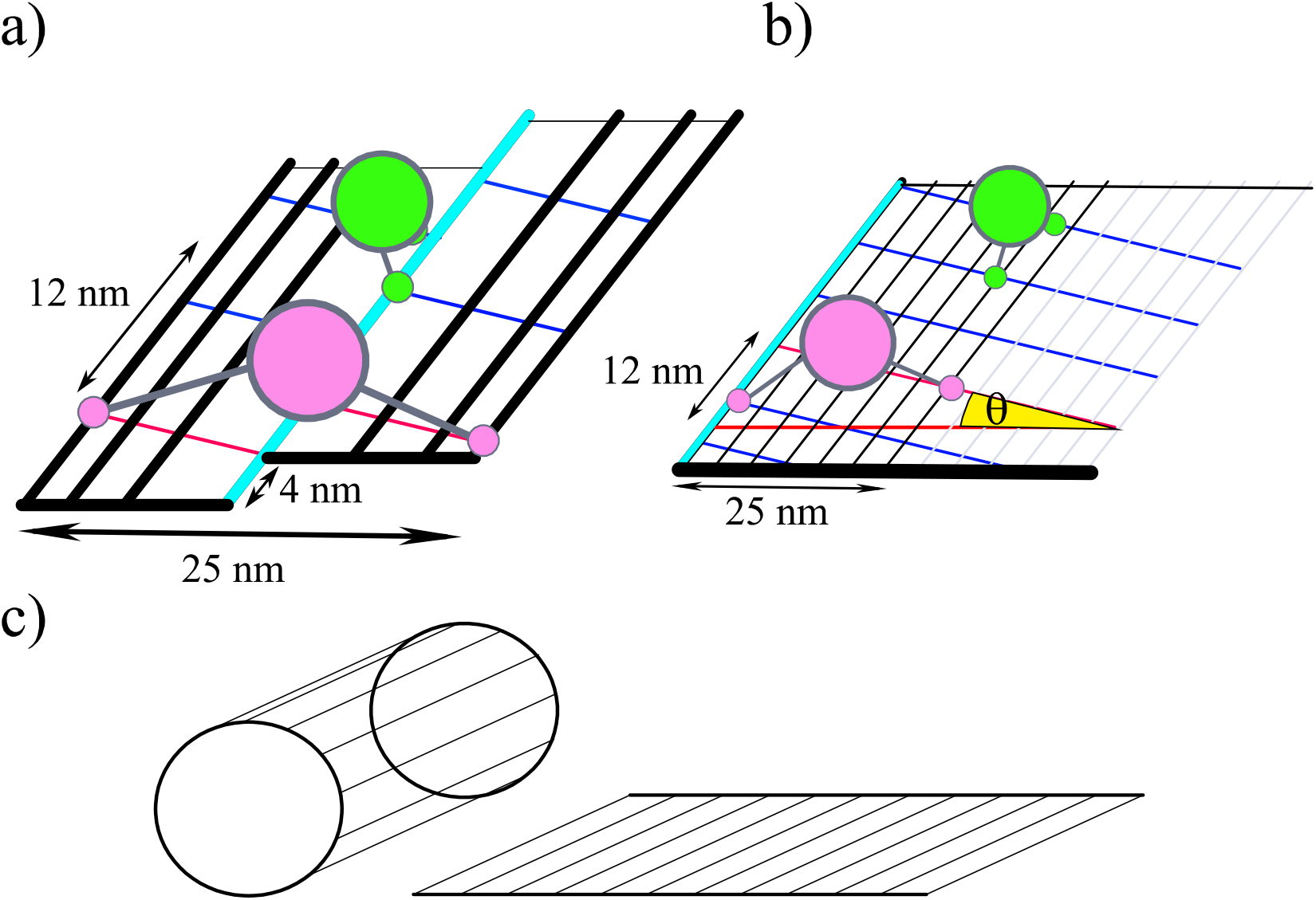
Illustration of the 2D array structure (in detail) and binding of dynein. Dynein motor protein pivots (pink and green big circles) and their MTBDs (pink and green small circles) binding to the MT lattice. Blue and red lines denote the transverse diagonals (note that the red line purpose is to emphasize the symmetric break, but it is not different from the blue lines). Black lines denote the protofilaments. Teal dashed line denote the seam that leads to a 4 nm longitudinal shift of the 2D binding array, leading to an angle *θ* (denoted in yellow) of 0.15 radians (or 7.5 degrees) between MT short axis (red dashed line) and the transverse diagonals (red and blue lines). Notably, for any perfect helix, the helix angle, *α* = π/2 − *θ*, must obey the following equation: *α* = tan^−1^(2*πr*/(*d* × *n*)) where r is the helix radius (12.5 nm for MT), *n* is an integer, *d* is the helical pitch (8nm for MT), and *d* × *n* is the helical rise after one encirclement (beginning at the helix edge). Since in MT case a non-integer value of *n* is required (*n* = 1.5), it is not a perfect helix and demands a seam defect. **(a)** The 2D binding array (as seen from above) with the seam at the middle, along with possible configurations for a MTBDs pair as they cross the seam. Notice that when MTBDs cross the seam, they break the rules and reside on the same diagonal but do not break the rules regarding the forbidden angles. **(b)** A different perspective of the same 2D array (unlike the case of **(a)**, here the MT cylnder is “spread” to a sheet), with the seam located at the left side, along with configurations of MTBDs that do not cross the seam.Thus, the MTBDs obey all the rules mentioned above without any exceptions. The pale gray lines indicate protfilaments that are not in the 25 nm width range. **(c)** Illustration of the spread of the MT 3D cylinder to a 2D sheet.

#### 2. The theoretical interplay between two MTBDs

Consider two MTBDs that belong to the same dynein molecule, henceforth referred to as MTBD1 and MTBD2. While the forces that act between them are fascinating *per se*, the explicit description of these forces is beyond the scope of this work. Instead, we chose to use experimental results as follows.

##### Angular distribution

According to the 2D attachment array (see Fig. 3), if hypothetically MTBD1 and MTBD2 have attached to the same diagonal and have not crossed the seam, the sharp angle between the separation vector (between the two MTBDs) and the *y*-axis would be 7.5 degrees. However, by marking the two AAA+ rings (“heads”) with different colors and measuring the angle between them (7), it has been shown that there is zero probability for 7.5 degrees angle between two rings (of the same motor). Although each marker is on the AAA+ ring and not on the MTBD itself (i.e., it is separated by a 15nm stalk, not precisely in the cylinder radial direction(2)), those measurements do give a strong indication regarding the MTBDs location. Thus, we assume that there is also zero probability for 7.5 degrees angle between the two MTBDs, suggesting that they cannot occupy the same diagonal, as long as they do not cross the seam. If the MTBDs do cross the seam, they can attach to the same diagonal without breaking the 7.5 angle limitation. Other than the latter angular constraint, we *do not use* the measured angular distribution explicitly.

##### Length distribution

Demanding that the two MTBDs do not occupy the same diagonal suggests that the shortest possible separation distance between them – the distance between two (parallel) neighboring diagonals – is 7.99 nm or ~4.62 nm if they cross the seam (see Fig. 3-a, b, Green colored motor and SI Figs. 1-c, 3). In addition, to account for the forces that act between MTBD1 and MTBD2, we use the probability histogram as presented in Qiu W, *et al*. work, Fig 5-b (We would like to underline the fact that numerical values of the inter-MTBDs distance probabilities were extracted from the figures in Ref. (7) with estimated error less than 10%, since we do not have the actual numerical data).

#### 3. A single theoretical MTBD translation vector probability distribution

The same experiment mentioned above (7) measures the angular distribution of the “heads” motion and shows that the “heads” never move in the 7.5 degrees’ direction. Using the same reasoning as in subsection 2, it follows that MTBD1 or MTBD2 never move in the 7.5 degrees’ direction. This implies that, unless the MTBD crosses the seam during its motion, MTBD cannot make a step along the same diagonal on which it already resides. As a result, considering the 25 nm array width (along the *y*-axis), |*ℓ_x_*| cannot be shorter than 4.25 nm (SI Fig.1-e) if the MTBD does not cross the seam, while it is about 0.37nm (SI Fig.3) if the MTBD does cross the seam, see Fig. 3-a,b. This implies that if the MTBD does not cross the seam, the smallest value of |*L_x_*| in a single MTBD stepping event is 2.125 nm (i.e., *L_x_* values between −2.125 to +2.125 nm are forbidden).

##### Stepping rules deduced from subsections 1-3

The discussion in subsections 1-3 leads to the following MTBD stepping rules. First and foremost, (i) each MTBD can only attach to specific sites on the MT surface. In addition, if the MTBDs do not cross the seam, (ii) the two MTBDs cannot reside on the same diagonal, and (iii) each one of them must change diagonal as it moves to another attachment site. As a consequence of rules (i)-(iii), as long as the MTBDs do not cross the seam, each MTBD cannot step onto a diagonal on which the other MTBD already resides. According to the above rules, the distribution for 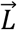 varies according to the temporal configuration of the two MTBDs.

#### 4. Inclusion of a longitudinal load force

An MTBD stepping probability function 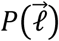 is taken as a product of two functions: *G*(*ℓ_x_*) and 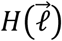. The former is the force exertion function (9), and the latter is a general function that accounts for the model rules that were mentioned above. *G*(*ℓ_x_*) accounts for the forward bias in terms of the motor internal force *F_s_* (working forward), which derives from ATP hydrolysis, and an external *longitudinal* pulling force *F*. It has been previously derived to be (9):

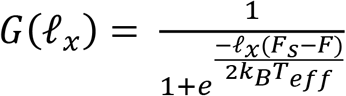

where *k_B_* is Boltzmann constant, and *T*_eff_ is an effective temperature that is taken as a fit parameter (see discussion at the end of the next section). By construction, the internal motor force is exactly equal in magnitude to the stall force, *F_s_*=7pN for yeast dynein (as described in Ref. (9)).

## Results

### Effective temperature of dynein

We applied the model rules 1-4 in Monte-Carlo computer simulations and computed the resulting distribution of 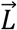 in ATP-affluent environment, with (or without) the exertion of external *longitudinal* pulling forces. When we use the room temperature (298°*K*) as *T*_eff_ to calculate the relative fractions of forward and backward steps in the case of ATP-affluent system and without an external force (*F* = 0), we find almost 100% of forward steps, which contradicts the experimental results reported in (7), 77% and 23%, of forward and backward steps, respectively. This apparent contradiction is resolved by replacing the actual temperature by the effective temperature that is used as a fit parameter. The effective temperature that recovers the experimental values mentioned above (77% and 23%, for forward and backward steps, respectively) is *T*_eff_ = 2540°*K*. The success of the use of a single effective temperature is further established by the excellent agreement (We assume that the seam is hardly sampled by the motor, so it does not affect the motor motion (Fig. 6)) between our model results and the experimental results reported in Ref. (3), in which dynein motion under different values of external pulling force was measured. Moreover, the mean *longitudinal* step of dynein that stems from *T*_eff_ = 2540°*K* is roughly 9 nm, while the use of room temperature leads to 12 nm. This value, 9 nm, is in agreement with several experiments that report a *longitudinal* step size distribution with a mode of 8 nm and a tail of long steps (> 8 nm) (7, 8).

### Dynein pivot mean stepping vector

The results for the angular distribution of the dynein pivot movement, under different longitudinal force values, are shown in Fig. 4. Note that although the angular distribution provides information regarding the relation between longitudinal- and side–steps, it does not represent the vector size of the dynein pivot. Therefore, for further analysis, we deduce the *L_x_* and *L_y_* distributions for two cases: in the first case, the dynein encounters the seam (Fig. 5), while in the second case, the dynein never encounters the seam (Fig. 6).

**Fig. 4.**
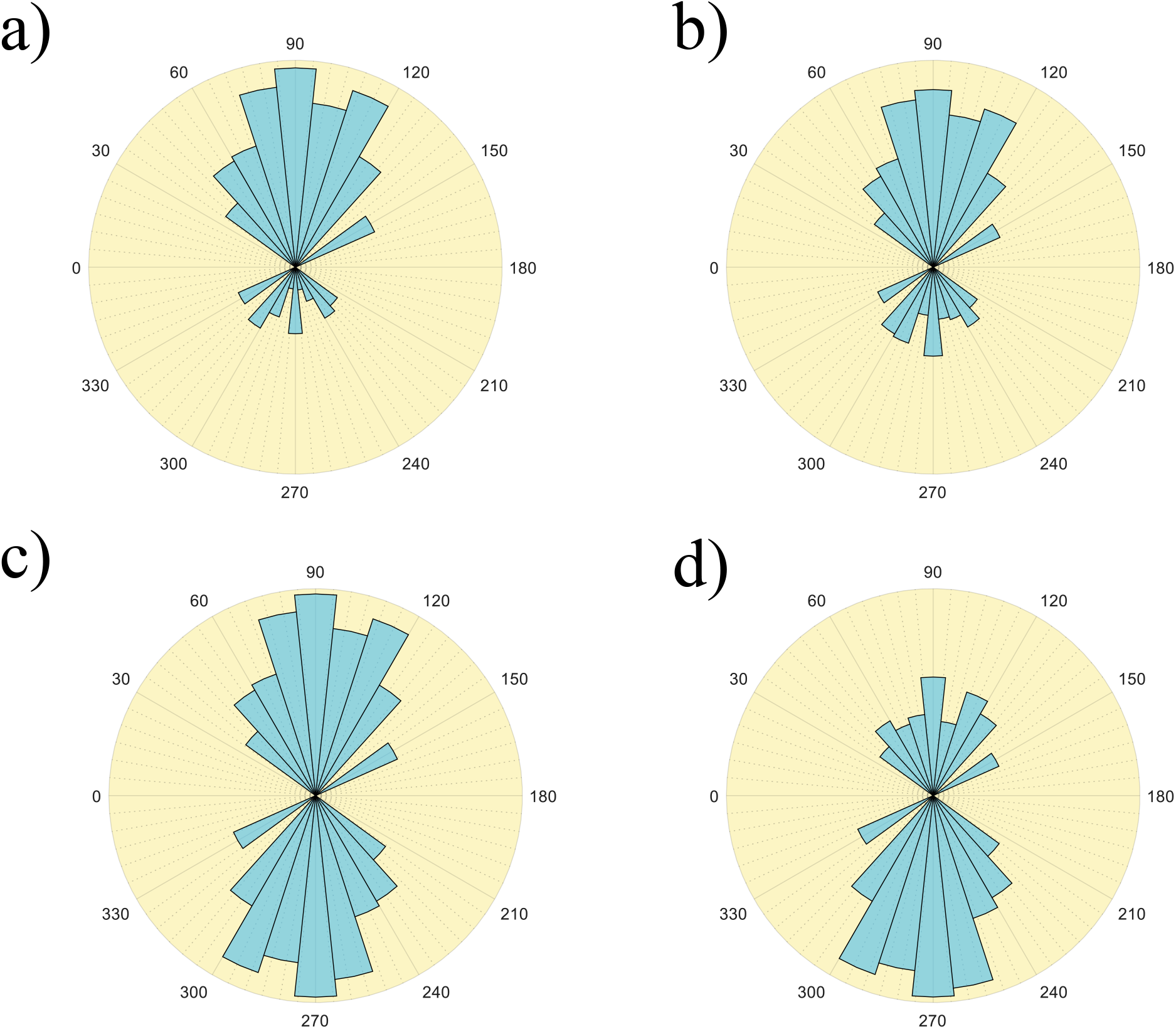
Angular histograms of MTBDs displacements under ATP-aflluent conditions and with an external force. The MT long axis is parallel to the 90°-270° axis (90° is the MT minus-end direction). The MT short axis is parallel to the 0°-180° axis; left and right (as referred to in the text) correspond to 0° and 180°, respectively. F is the longitudinal pulling force, meaning that it pulls against the motor direction towards the MT plus-end **(a-d)** Angular histogram for F = 0,3,7 and 10 pN with a mean angle of 139 ± 0.0171, 154 ± 0.0171, 180 ± 0.0171 (or 0), 205 ± 0.0171, respectively (SEM, N = 25,000,000 each).

**Fig. 5.**
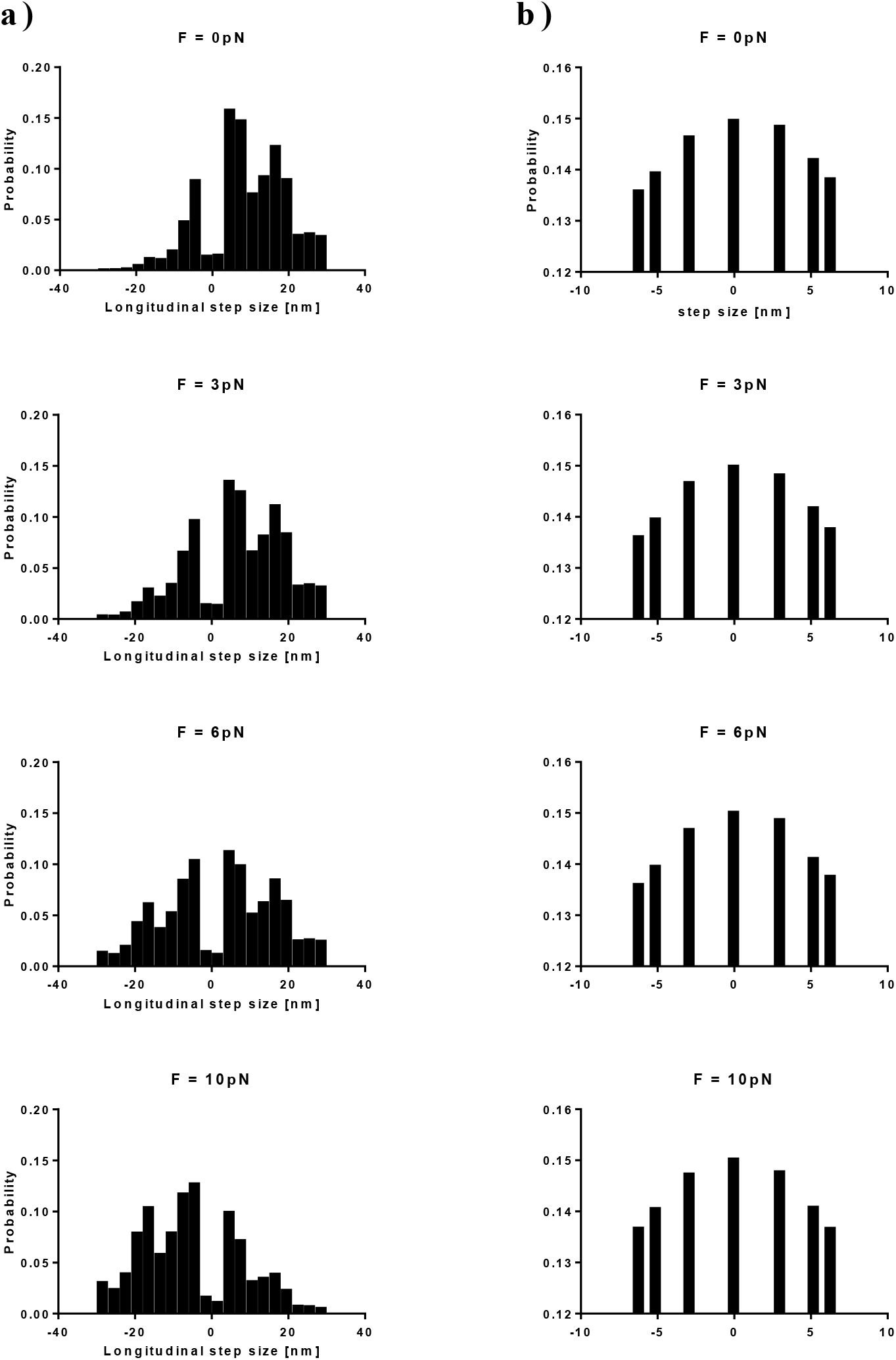
Dynein pivot displacements under ATP-affluent conditions and under exertion of a pulling longitudinal force, for a pseudo-helical MT structure (note that our simulations length leads adequate sampling of the seam) Histograms computed from 2.5 × 10^7^ computer steps each. **(a)** Displacements along the MT *x*-axis (longitudinal steps). **(b)** Displacements along the MT *y*-axis (sidesteps).

**Fig. 6.**
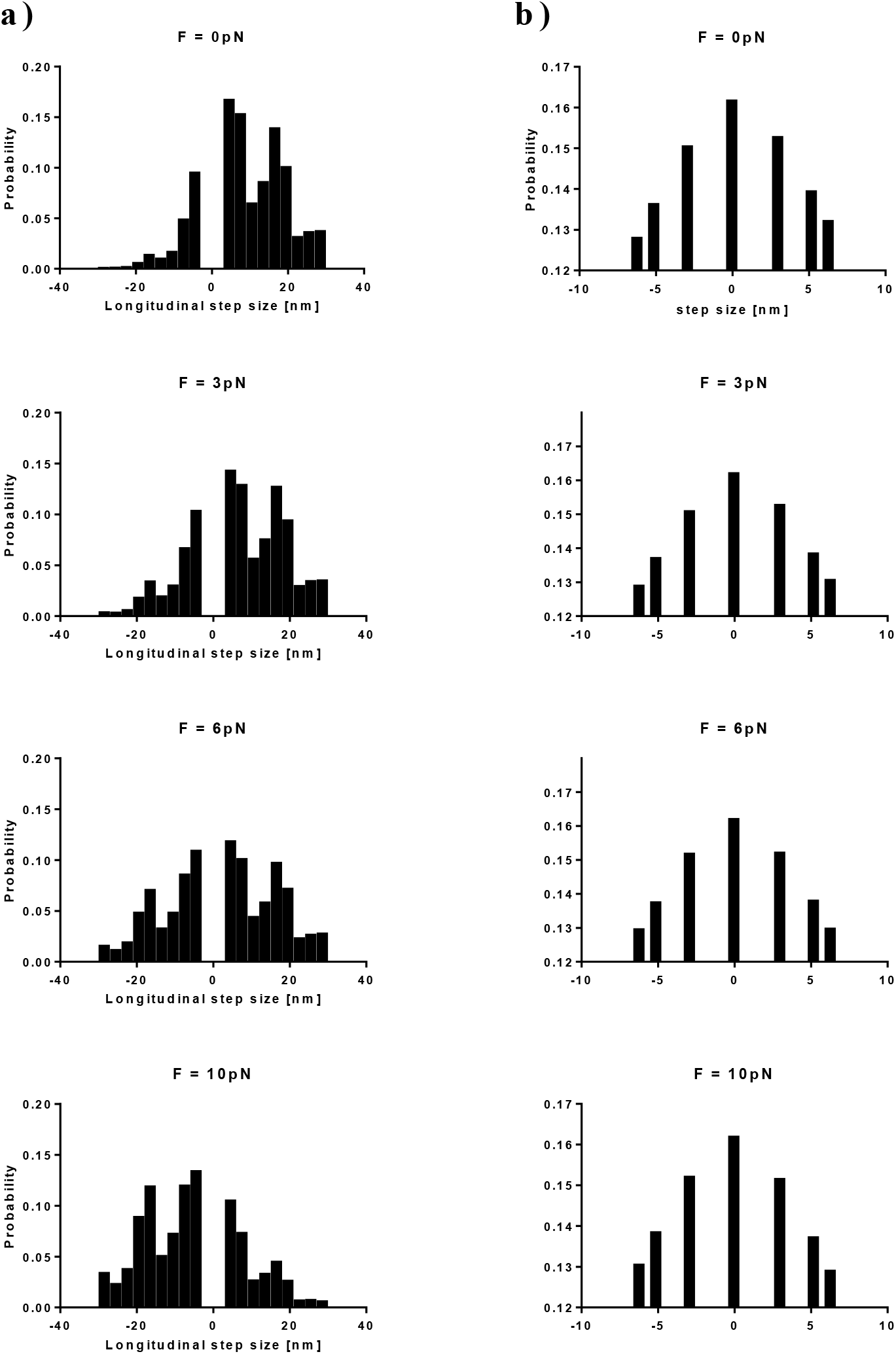
Same as Fig 5 but for the unique case of zero encounters with the seam line. I.e, the MTBDs never cross the seam line (or does not sample the seam enough times for its effect to be discerened).

As expected, the *longitudinal* (*L_x_*) distribution++n in Figs. 5-a and 6-a show that when *F* = 0 pN, forward steps (*L_x_* > 0) prevail, when *F* = *F_s_*, the distribution becomes symmetric, and when *F* > *F_s_*, the distribution is again asymmetric with a predominance of backward steps (*L_x_* < 0). Fig. 6-a shows a “hole” in the *L_x_* distribution around the origin, between −3.065 *nm* < *L_x_* < +3.065 nm, which is related to stepping rule (iii) and the previous discussion in subsection 3. On the other hand, in Fig. 5-a, where the dynein encounters the seam, the “hole” is absent.

The *transverse* (*L_y_*) distribution, depicted in Figs. 5-b and 6-b, indicates that the motion of single dynein in its native state (*F_s_* = 7pN, *F* = 0) is slightly biased towards the right-hand side of the *y*-axis (*L_y_* > 0). Calculating the mean *transverse* component from the distribution leads to < *L_y_* > ≅ 0.048 ± 0.0028 nm (SEM; N = 2.5 × 10^7^). As the pulling force increases from zero, the distribution becomes less biased and achieves complete symmetry for *F* = *F_s_*, as the distribution for *L_x_*. When *F* > *F_s_*, the bias shifts towards the left hand side of the *y*-axis (*L_y_* < 0). Moreover, as the pulling force decreases from zero and turns into an assisting force, the mean *transverse* component value significantly increases (see Fig. 7), until it reaches a plateau for large enough forces (see SI Fig 4). Note that the exerted force is purely *longitudinal*, nevertheless, it still affects the *transverse* motion, implying that the cross Onsager coefficients of linear response of the motor to force, coupling the *x* and *y* axes, are non-zero (23).

**Fig. 7.**
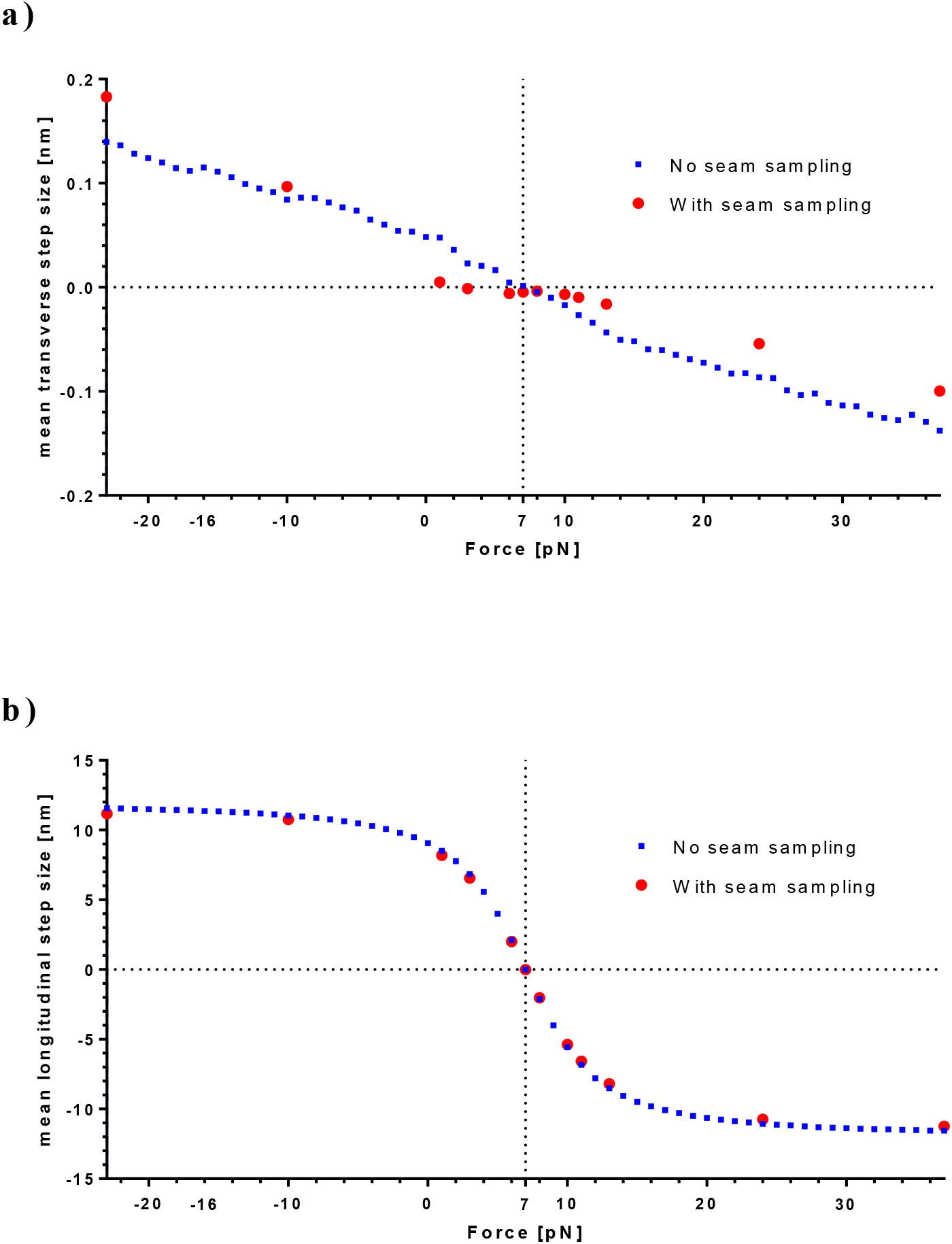
Force-dependence of longitudinal and transverse mean step size. The mean longitudinal 〈*L_x_*〉 and transverse step 〈*L_y_*〉 against longitudinal pulling force *F*; a negative force indicates an assisting force (i.e., a force that is in the MT minus-end direction), and positive indicates a resisting force (i.e., a force that is in the MT plus-end direction). Red and blue dots denote *F* = 7 and *F* =10 pN, respectively. Note, if one assumes equal dwell time *τ* for all steps, independent of the exerted force, the motor velocity is given by *v* = 〈*l*〉/*τ*. Therefore, these graphs also depict the velocity trends. **(a)** Transverse mean step size against force (SEM = 0.0021 nm, 2.5 × 10^7^ computer step for each force value). We note that regarding the results with the seam being sampled (red dots), the statistical error is likely larger than the above numerically estimated value due to insufficient number of seam sampling events. **(b)** Longitudinal mean step size against force (SEM = 8.98 × 10^−4^ nm, 2.5 × 10^7^ computer step for each force value).

## Discussion and Conclusions

In this work, we presented a model that aims to predict yeast dynein motion on the 2D microtubule surface. Our model assumptions are based on several distinct and independent experimental results. First, we considered data that implies the existence of specific MTBD attachment sites on the MT surface. Second, we incorporated the interplay between a pair of MTBDs, i.e., the distance and angle probability distributions between two MTBDs.

The present study can be regarded as a first principle calculation of the (phenomenological) bare dynein motor step size distribution under stall force conditions (denoted as *f*_0_(*ℓ*) in (9)). Moreover, our model predictions for a system under ATP-affluent conditions and exertion of external *longitudinal* pulling force are consistent with a variety of experimental results. Notice that we did not examine the case of ATP-depleted system since experimental results indicate that dynein remains stagnant under such conditions (3).

Let us discuss first the *L_x_* distribution (as depicted in Figs 5 and 6). Under external force, our model predictions are as per those described in (3). Also, our model suggests a plausible explanation for the “hole” (i.e., vanishing probability) around the origin in the *L_x_* distribution, as measured in a few experiments (3, 8, 10, 11). Yet, according to our model, the effective size of the “hole” depends on whether the seam has been encountered or not. Therefore, we believe that we have acquired a reasonable explanation for the seeming discrimination between different experiments regarding the magnitude of the “hole,” e.g., a deep and wide “hole” or a smeared one.

Next, consider dynein *transverse* motion (*L_y_*) and the effect of external longitudinal pulling forces over such motion. Under native conditions and without external force (*F_s_* = 7pN, F = 0), regardless if the seam is adequately sampled or not sampled at all, the *L_y_* distribution shows a meager bias towards right side-steps. For the case in which the seam is not sampled adequately, a situation we assume to prevail in most cases, < *L_y_* ≅ = 0.048 ± 0.0028 nm (SEM; N = 2.5 × 10^7^)). Thus, combining this result with the prediction for the mean longitudinal step size in the absence of external force, < *L_x_* >= 9 ± 0.0066 nm (SEM; N = 2.5 × 10^7^), a single motor is predicted to move in a right-handed helical pattern with helical pitch size of 14.7 μm. Obviously, such a motion cannot be experimentally observed, since the required longitudinal run length of at least 14.7μm exceeds the mean run length of 1.9 μm (8) by more than sevenfold. Thus, we believe that, in the absence of external force, the *transverse* motion might usually *seem unbiased*. Surprisingly, we also found that when the *longitudinal* pulling force (*F*) is moderately larger than the stall force (*F_s_*), the *L_y_* distribution shifts to give a small bias towards left-handed side-steps (*L_y_* < 0), describing a motor that is walking backward and making a right-handed *helical* motion. While a small or moderate longitudinal force does not result in significant bias, the exertion of an adequate *longitudinal* force (whether it is pulling or assisting), enhances the transverse bias significantly (see Fig. 7-b), e.g., for (an assisting) force *F* = −10pW, < *L_y_* >≅ 0.084 ± 0.0028 nm (SEM; N = 2.5 × 10^7^), corresponding to a right handed helix with a pitch size of 8.4 μm. Measuring the side-steps distribution under different values of longitudinal pulling force would thus be highly valuable to confirm these non-trivial predictions of our model. Although the exertion of a *transverse* force is also likely to enhance the *L_y_* bias, we did not include such forces due to lack of experimental results that could be used as validation.

Importantly, measurements of the helical motion of a bead, possibly being carried by more than one motor, have been reported in experiments in which the MT is lifted (well above the flow chamber surface), thus allowing free motion of the motors on the MT surface (5). The agreement is however only partial. First, the observed pitch helical size is ~500-600 nm, much shorter than our prediction. Second, S. Can *et al*. also observed left-handed helical motion. We believe that these differences stem from the fact that very likely (as also mentioned by the authors of Ref. (5)) several motors are carrying the particle simultaneously. Therefore, inter-motor coupling forces, which include both *longitudinal* and *transverse* components, may strongly affect the motion. This ought to motivate further investigations.

What is the physics behind the effective temperature? We believe that it is caused by the *partial* efficiency in converting the chemical energy, released in ATP-hydrolysis, to mechanical work. This implies that part of the stored energy renders into an active force noise, unable to instantaneously dissipate into the surrounding medium. Active noise has been indeed shown in a variety of active systems to be equivalent to an effective temperature that can be several times higher than room temperature. Indeed, since a single ATP hydrolysis releases about 8.3 × 10^−20^J = 20 *k_B_T* (24), it follows that for *T*_eff_ = 2540 *K* the efficiency, estimated from the equality *k_B_T*_eff_ = (1 − *η*)20 *k_B_T*, is about *η* = 58%, which is well within the range of reasonable values for an efficient molecular machine.

What can our results imply regarding the motion of mammalian cytoplasmic dynein? Our model can be extended to describe this motor by assuming similar MT-MTBD interactions, however with two modifications. First, the *F_s_* value must be changed to 1.1pN – the mammalian stall force (4). Second, the MTBDs separation vector distribution of mammalian dynein must be acquired. We hope that investigations concerning the latter will be carried out in the future.

## Supporting information

Model details and additional results

## Author contributions

G.R. & F.I. designed the model, analyzed data, and wrote the manuscript.

## Acknowledgements

We thank Anne Bernheim for useful and illuminating discussions.

