## Supplementary material for "Directional stepping model for yeast dynein: Longitudinal- and side- step distributions": Model details and additional results

### Supplementary Information

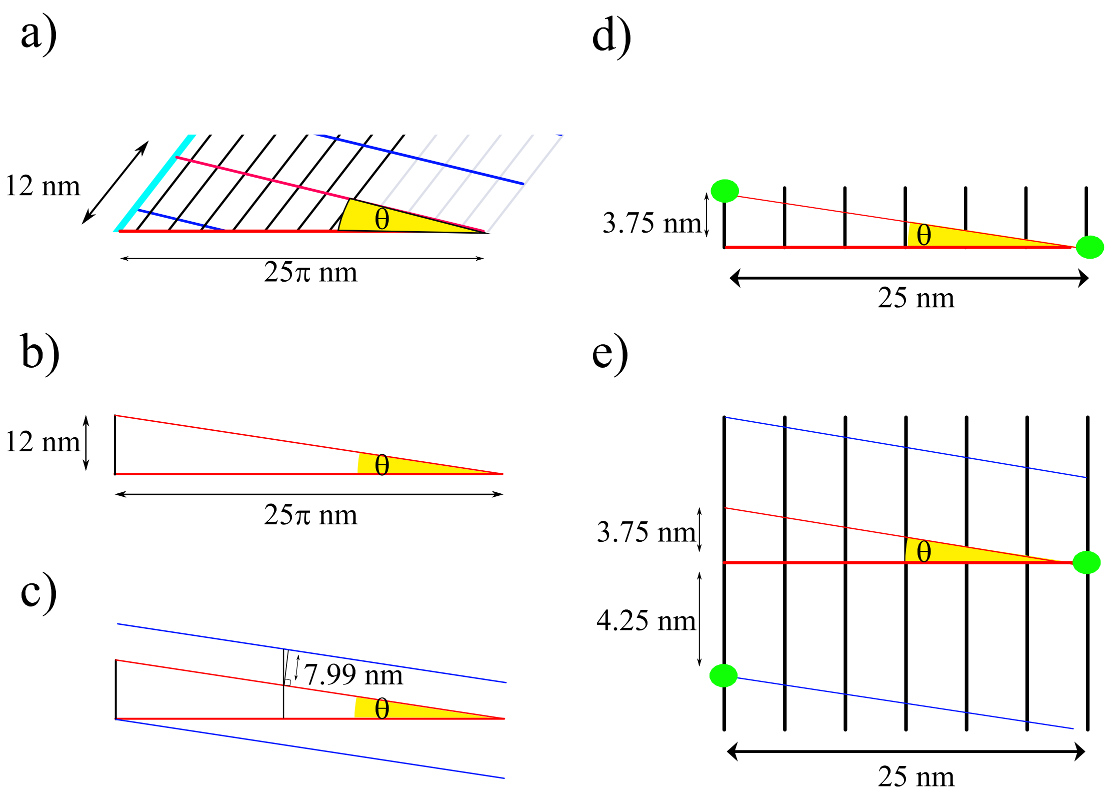


**S.I Fig. 1 – Calculation of the maximum distances if the MTBDs do not cross the seam:** Here we show that, (i) as long as the longitudinal distance between two MTBDs is longer than 4.25 nm, the MTBDs reside on different transverse diagonals, and (ii) as long as the value of a single MTBD longitudinal component is larger than 3.75 nm, the MTBD does not step to the same diagonal on which it already resides. In (a)-(e), the seam is denoted by teal dashed line, MTBDs location denoted by green dotes, transverse diagonals by red and blue lines, tubulin protofilaments by black lines, and MT short axis by red dashed line. **(a)** Enlargement of Fig. 3-b. **(b)** Calculation of $\theta$ between the MT short axis and the transverse diagonals. The first leg of the right angle triangle is the helical rise; we take it to be 12 nm, the sum of the distance between adjacent transverse diagonals (8 nm) and the seam shift (4 nm). The second leg is the helix (or cylinder) circumscribe. $\theta = \tan^{-1}(12/25\pi) \cong \mathbf{0.15}$ (radians) **(c)** Illustration of the shortest distance between two adjacent transverse diagonals. $8 * \cos\theta = \mathbf{7.99}$ nm. **(d)** The longest longitudinal distance between two MTBDs, if they could reside on the same diagonal, located within the array width of 25 nm. $25 \times \tan\theta = \mathbf{3.75}$ nm. **(e)** The shortest longitudinal distance between two MTBDs, if they reside on distinct, but adjacent, diagonals. **4.25** nm.

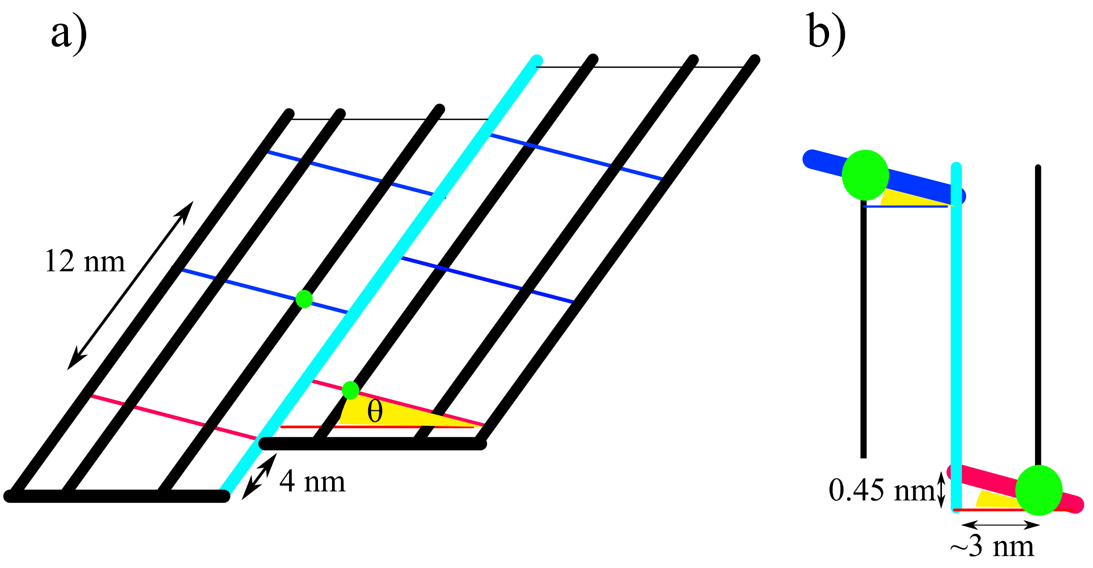


**S.I Fig.2 – Calculation of maximum distances for the case in which the MTBDs cross the seam.** The seam is denoted by teal dashed line, MTBDs location – by green dotes, transverse diagonals – by red and blue lines, tubulin protofilaments by black lines, and MT short axis by red dashed line. **(a)** Enlargement of Fig. 3-a. **(b)** Obviously, MTBDs do not bind precisely on the seam but in a certain distance from it. If we split the MT circumference to 13 segments (with respect to 13 protofilaments), each protofilament occupy an arc segment A of length $\frac{25\pi}{13} \cong$ **6** nm. Therefore, the shortest distance between the binding site and the seam is about 3 nm, resulting in a minimum distance of **4.62** nm between MTBDs (using the relation between longitudinal and transverse coordinates of the binding sites as defined by $\theta$).

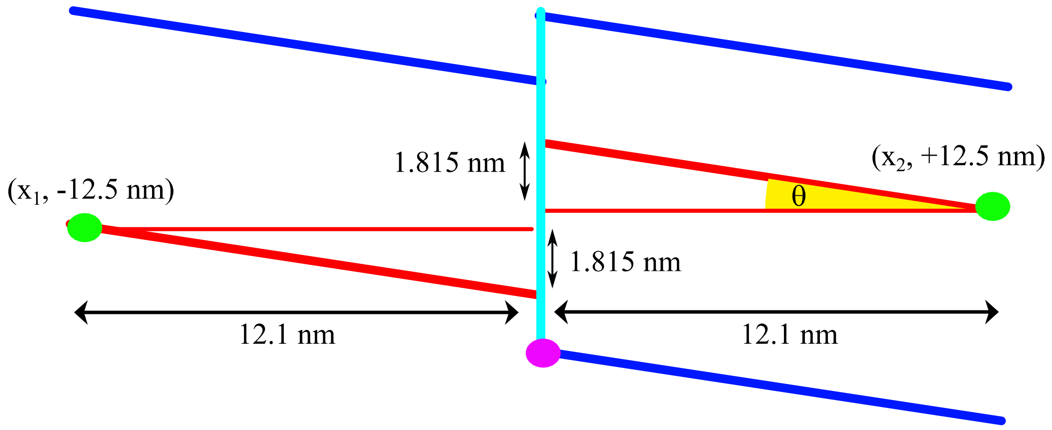


**S.I Fig.3 – Calculation of maximum longitudinal distance for the case in which the MTBDs cross the seam.** The seam denoted by teal dashed line, MTBDs location by green dotes, transverse diagonals by red and blue lines, tubulin protofilaments by black lines, and MT short axis by red dashed line. In this illustration, we demonstrate a step of a single MTBD (green) while the other MTBD (pink) stays static. The seam that is located in the middle is used as a reference point. The "before" and "after" step binding sites (green dots) with the coordinates

$(x_1, -12.5)$ and $(x_2, 12.5)$ nm lead to a longitudinal step size ($= |x_1 - x_2|$) of **0.37** nm (x1 and x2 are calculated through the relation between the x and y coordinates as defined by the angle $\theta$ )

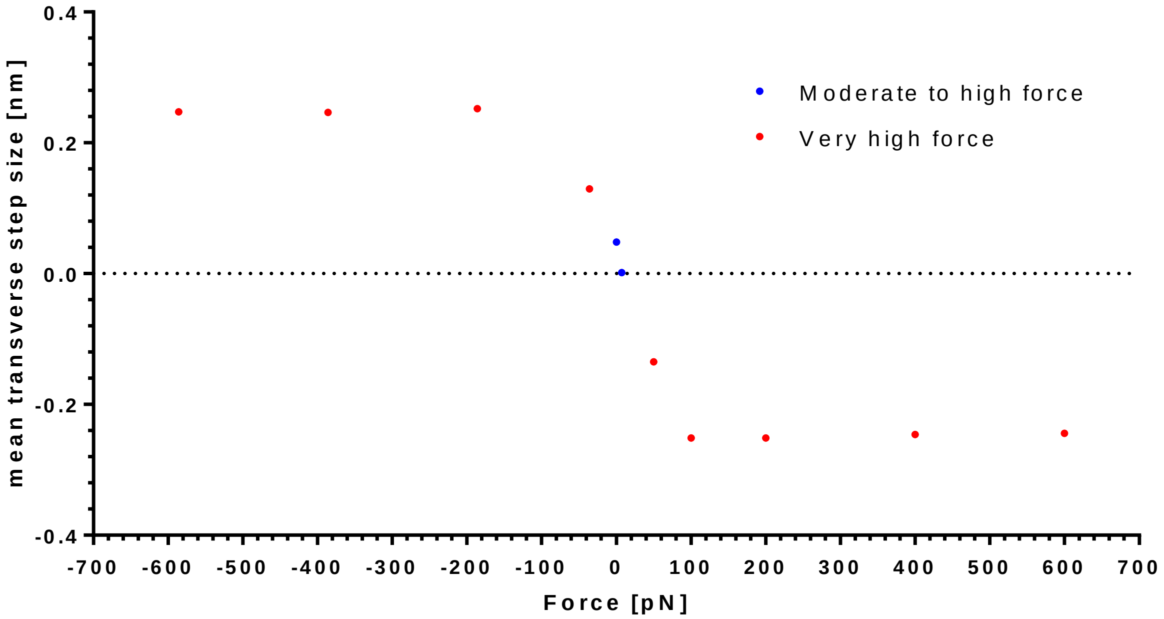


**S.I Fig.4 - Transverse mean step against very high forces.** Blue dots denote selected moderate forces, which are used as reference points, and red dots denote very high forces. For pulling forces, i.e., $\gg F_s$ , the mean transverse reaches a plateau value of $< L_y > \cong -0.24\, nm$; for assisting forces, i.e., $F \ll F_s$ , the mean transverse reaches a plateau value of $< L_y > \cong +0.24\, nm$.
